# CRISPR/Cas9-based silencing of the *ATXN1* gene in Spinocerebellar ataxia type 1 (SCA1) fibroblasts

**DOI:** 10.1101/2020.07.04.187559

**Authors:** Francesca Salvatori, Mariangela Pappadà, Mariaconcetta Sicurella, Mattia Buratto, Valentina Simioni, Valeria Tugnoli, Peggy Marconi

## Abstract

Spinocerebellar Ataxia type 1 (SCA1) is an autosomal dominant neurodegenerative disorder caused by a gain-of-function protein with toxic activities, containing an expanded polyQ tract in the coding region. Actually, there are no treatments available to delay the onset, stop or slow down the progression of this pathology. Many approaches developed over the years involve the use of siRNAs and antisense oligonucleotides (ASOs). Here we develop and validate a CRISPR/Cas9 therapeutic strategy in fibroblasts isolated from SCA1 patients. We started from the screening of 10 different sgRNAs able to recognize regions upstream and downstream the CAG repeats, in exon 8 of *ATXN1* gene. The two most promising sgRNAs, G3 and G8, whose efficiency was evaluated with an *in vitro* system, significantly downregulated the ATXN 1 protein expression. This downregulation was due to the introduction of indels mutations into the *ATXN1* gene. Notably, with an RNA-seq analysis, we demonstrated minimal off-target effects of our sgRNAs. These preliminary results support CRISPR/Cas9 as a promising approach for treated polyQ-expanded diseases.

## Introduction

SCA1 is an autosomal dominant neurodegenerative disorder caused by a CAG-repeat expansion in *ATXN1* gene. The severity of the disease and its onset are directly proportional to the number of triplets. The mutated ATXN1 contains an expanded polyQ tract and shows new toxic functions which lead to neurodegeneration [Orr, 2000; Zoghbi and Orr, 2009]. PolyQ SCAs, which have a frequency of 2-3 cases per 100.000 people, are progressive, typically striking in midlife and causing increasing neuronal dysfunction and eventual neuronal loss 10-20 years after onset of symptoms [Zoghbi and Orr, 2000]. Patients can lose the ability to breathe in a coordinated fashion, which can be fatal [Orr, 2012].

Currently, no treatments are available to prevent or cure, delay the onset, stop or slow down the progression of these diseases. Several research groups are trying to develop strategies to reduce the levels of mutated proteins, and therefore their neurotoxic effects, using for example antiaggregant agents [Zoghbi and Orr, 2000], molecules capable of activating the ubiquitin-proteasome pathway [Nagashima et al., 2011] and compound with autophagic effects, as lithium [Watase et al., 2007], tensirolimus [Menzies et al., 2010], thehalose [Chen et al., 2015] and Beclin-1 [Nascimento-Ferreira et al., 2011].

Potential genetic therapies for SCA1 involve several different nucleic acid-based molecules able to target the RNA or DNA of the polyQ-associated genes. Since methods exploiting the mechanisms involved in the processing of endogenous miRNAs, as siRNAs and shRNAs, have shown potential toxicity [Grimm et al., 2010], antisense oligonucleotide-mediated (ASO-mediated) RNA suppression approaches have been recently used to reduce gene expression and improve disease symptoms in preclinical rodent models of several neurological diseases [Friedrich et al., 2018; McLoughlin et al., 2018], including SCA1 [Friedrich et al., 2018]. The disadvantage of these methods lies in the need for continuous administration throughout the patient’s life, to keep toxic protein levels low.

To address this fundamental limitation, the field of gene editing has emerged to make precise, targeted modifications to genome sequences. The most popular and used tool for gene editing currently is the clustered regularly interspaced short palindromic repeat (CRISPR)/Cas9 system [Jinek et al., 2012]. The CRISPR/Cas9, a component of the bacterial RNA-mediated adaptive immune system, consists of transcribed guide RNAs that direct the Cas9 RNA-guided DNA endonuclease to target sequences [Jinek et al., 2012; Barrangou et al., 2007]. The CRISPR/Cas9 system from *Streptococcus pyogenes* has already been successfully used in treating genetic disorders [Jinek et al., 2012; Jinek et al., 2013].

## Results

### Design and *in vitro* screening of sgRNAs

We designed ten different sgRNAs able to recognize sequences upstream and downstream the *ATXN1* polyQ tract (Fig. 1A), using the CRISPR design software developed by Zhang lab [Brezelton et al., 2015], and tested the efficiency to recognise and mediate the cutting by Cas9 of these sgRNAs in *in vitro* reactions. The efficiencies shown in figure 1b were determined as a percentage of the fragments obtained from the Cas9 cut related to the total amount of the target sequence. At least, four sgRNAs show a high capacity to mediate the cut of the target sequence (G3, G8, G10, G11), two of which recognize sequences upstream and two downstream of the polyQ tract. Since a multiple cut of the *ATXN1* gene can determine loss of genetic material and therefore a more efficient silencing of the gene itself [Maeder and Gersbach, 2016], we set up four pairs of sgRNAs, whose recognition and cutting efficiency was always very high (Fig. 1C). From preliminary tests on U266 cellular line, we chose the G3-G8 sgRNA pair which confirmed the ability to effectively knock-out *ATXN1* gene expression (data not shown).

**Figure 1.**
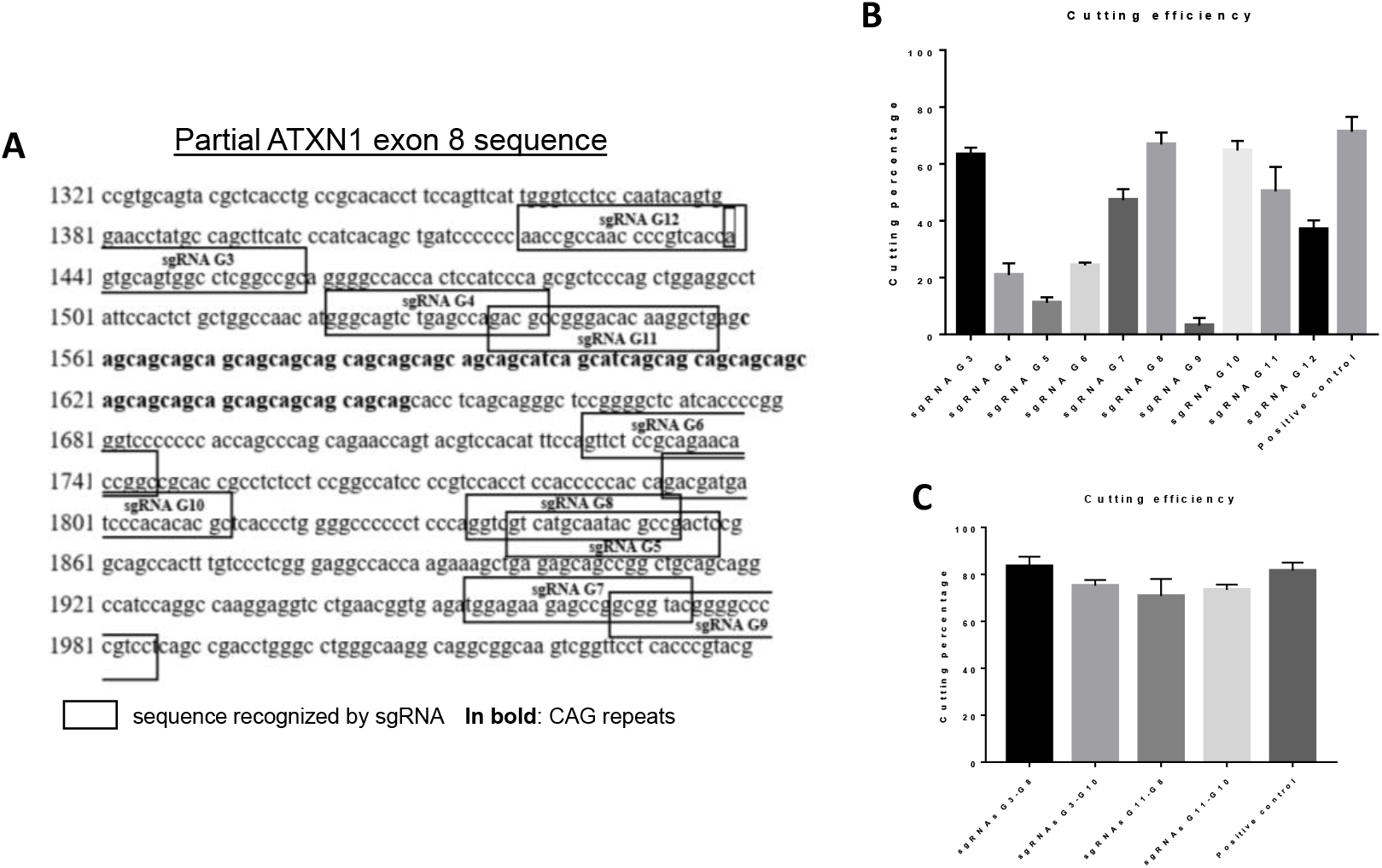
Design and *in vitro* screening of sgRNAs for ATXN1 gene. **A,** sgRNAs were designed using the CRISPR design software by Zhang lab. Ten sgRNAs capable of cutting upstream and downstream of the polyQ tract were identified. **B,** Using Guide-it sgRNA *In Vitro* Transcription and Screening System (Clontech), i*n vitro* screening tests were performed to evaluate the sgRNAs cutting efficiency, which were 63,3 ± 1,4 (G3), 20,9 ± 2,4 (G4), 11,1 ± 1,1 (G5), 24,3 ± 0,6 (G6), 47,3 ± 2,2 (G7), 66,8 ± 2,4 (G8), 3,2 ± 1,5 (G9), 64,7 ± 2,0 (G10), 50,4 ± 4,9 (G11), 37,0 ± 1,8 (G12). **C,** The most efficient sgRNAs were then tested in pairs. The cutting efficiencies were 83,4 ± 2,4 (G3-G8), 75,2 ± 1,4 (G3-G10), 70,8 ± 4,2 (G11-G8), 73,4 ± 1,3 (G11-G10). Values are mean ± s.e.m. from at least three independent experiments.

### Isolation and characterization of SCA1 fibroblasts

Fibroblasts were isolated from a 4 mm skin fragment, obtained from five patients, after the signature of the informed consent, with different polyQ expansion in mutated *ATXN1* gene: SCA1N1 (50 repeats), SCA1N5 (45), SCA1N6 (42), SCA1N8 (60), SCA1N17 (67).

To confirm the presence of an *ATXN1* allele with pathological increase in CAG triplets we amplified by PCR the exon 8 of the *ATXN1* gene and verified, by electrophoresis on agarose gel, the presence of two PCR fragments, one due to the healthy allele and one to the mutated one (Fig. 2a). At the same time, to verify the production of the wild type and expanded polyQ ataxin 1, a western blotting experiment was set up, where using anti-ATXN1 antiserum 12NQ it was possible to highlight the presence of both ATXN1 proteins (Fig. 2a).

**Figure 2.**
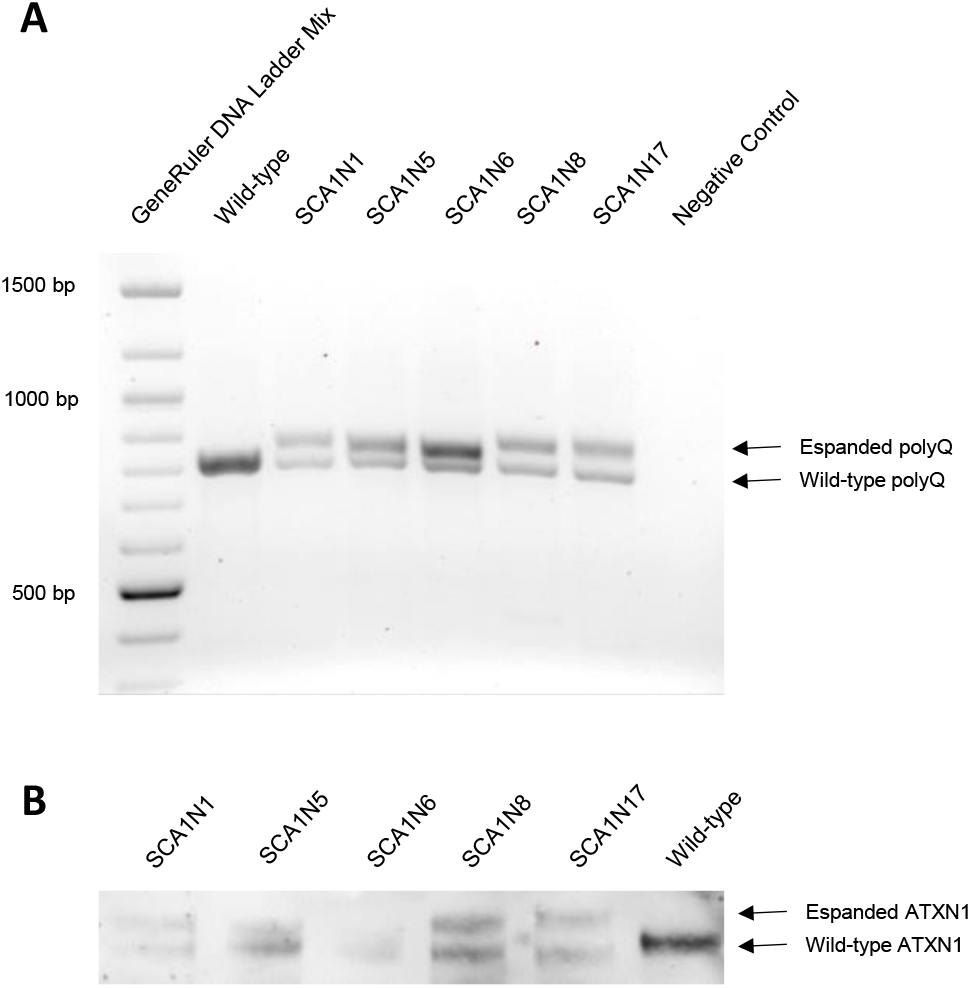
Characterization of SCA1 fibroblasts. **A**, Amplification by PCR of the *ATXN1* exon 8 containing the polyQ tract, using the genomic DNA extracted from the five SCA1 patients’ fibroblasts as template. **A**, ATXN1 expression in SCA1 fibroblasts, using 12NQ antiserum as anti-ATXN1 antibody.

### Validation of CRISPR/Cas9-based therapeutic strategy in SCA1 fibroblasts

Fibroblasts were then transfected with G3 and G8 sgRNA simultaneously and with a negative control sgRNA, as nucleofection control, previously associated with a Cas9 to obtain ribonucleoproteins (RNPs). Western blotting analysis with anti-ATXN1 antiserum 12NQ, using total protein amount and anti-HSP90 for normalization, showed significant reduction of the ATXN1 expression (from 27,2% to 75,2% of expression compared to not treated cells) (Fig. 3a,b).

**Figure 3.**
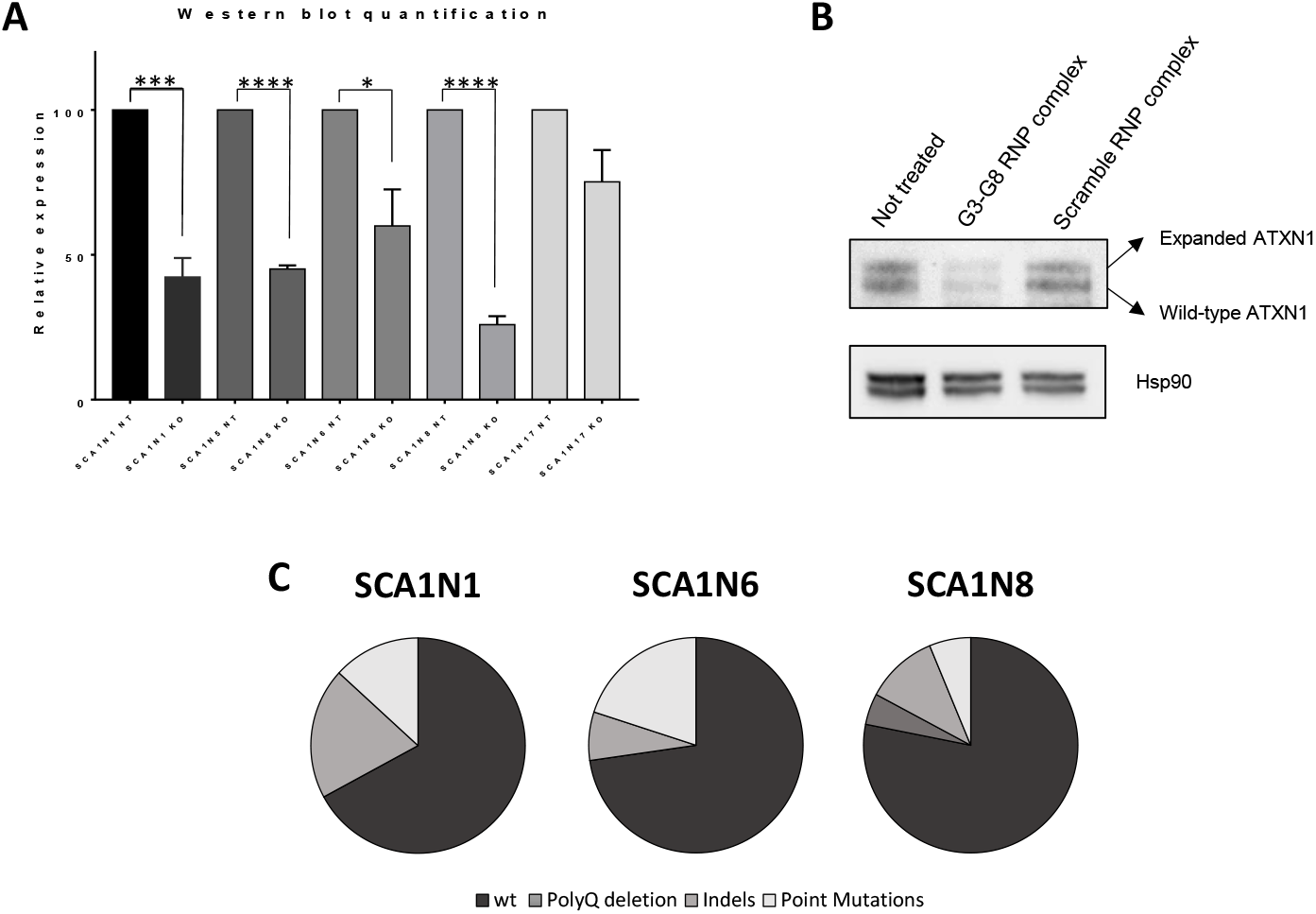
Effects of CRISPR/Cas9 system in SCA1 fibroblasts. **A-B**, Atxn1 expression in SCA1 fibroblast. Fibroblasts from five SCA1 patients were treated using sgRNAs G3 and G8 complexed with Cas9 endonucleases and the Atxn1 expression was determined by Western Blotting. **A**, Atxn1 abundances were expressed relative to Hsp90 and total proteins, determined by densitometry. After the treatment, the Atxn1 expression was 42,3 ± 3,8 (SCA1N1), 46 ± 2,6 (SCA1N5), 60 ± 5 (SCA1N6), 23,7 ± 2,6 (SCA1N8) and 75,2 ± 4,8 (SCA1N17). **B**, Expression in patient SCA1N1 for Atxn1 and HSP90. **C-E**, Mutations introduced by CRISPR/Cas9 system in ATXN1 genes of three patients and their relative abundance, determined by sequencing 76 (**C**), 55 (**D**) and 64 (**E**) colonies. Values are mean ± s.e.m. from at least three independent experiments. *P=0,05, ***P<0,001, ****P<0,0001, unpaired t test followed by f test to compare variances.

To verify which genetic alterations were introduced by our CRISPR/Cas9 system, we decided to amplify the region of exon 8 containing the polyQ and the sequences recognized by G3 and G8 of the treated cells of three patients and to subclone PCR products. Following sequencing and alignments with the correct exon sequence, we were able to determine in what percentage indels and point mutations were introduced or the polyQ was completely excised (Fig. 3c-e), obtaining 13% ± 4% for indels, 14% ± 5% for point mutations and 2% ± 2% for polyQ deletions.

### Evaluation of off-target effects

To verify if the G3 and G8 Cas9/sgRNA complexes determined a significant alteration in the expression of off-target genes, we carried out transcriptome profiling of the treated versus not treated fibroblasts of three patients by RNA-seq, performed by Lexogen. From a minimum of 5.44 to a maximum of 6.9 million uniquely mapped reads were carried out, for a total of approximately 1600 genes. Figure 4a displays inter-replicate correlations plots. The overall correlation of expression between samples which were defined as replicates is very good.

Differential expression analysis (Fig. 4b) was normalised using the DESeq methods [Anders and Huber, 2010], which ignore highly variable and/or highly expressed features. From this analysis it emerged that only two genes are significantly altered by treatment with Cas9/sgRNA complexes: *TXNIP* up regulated and *HAS2* down regulated about twice.

**Figure 4.**
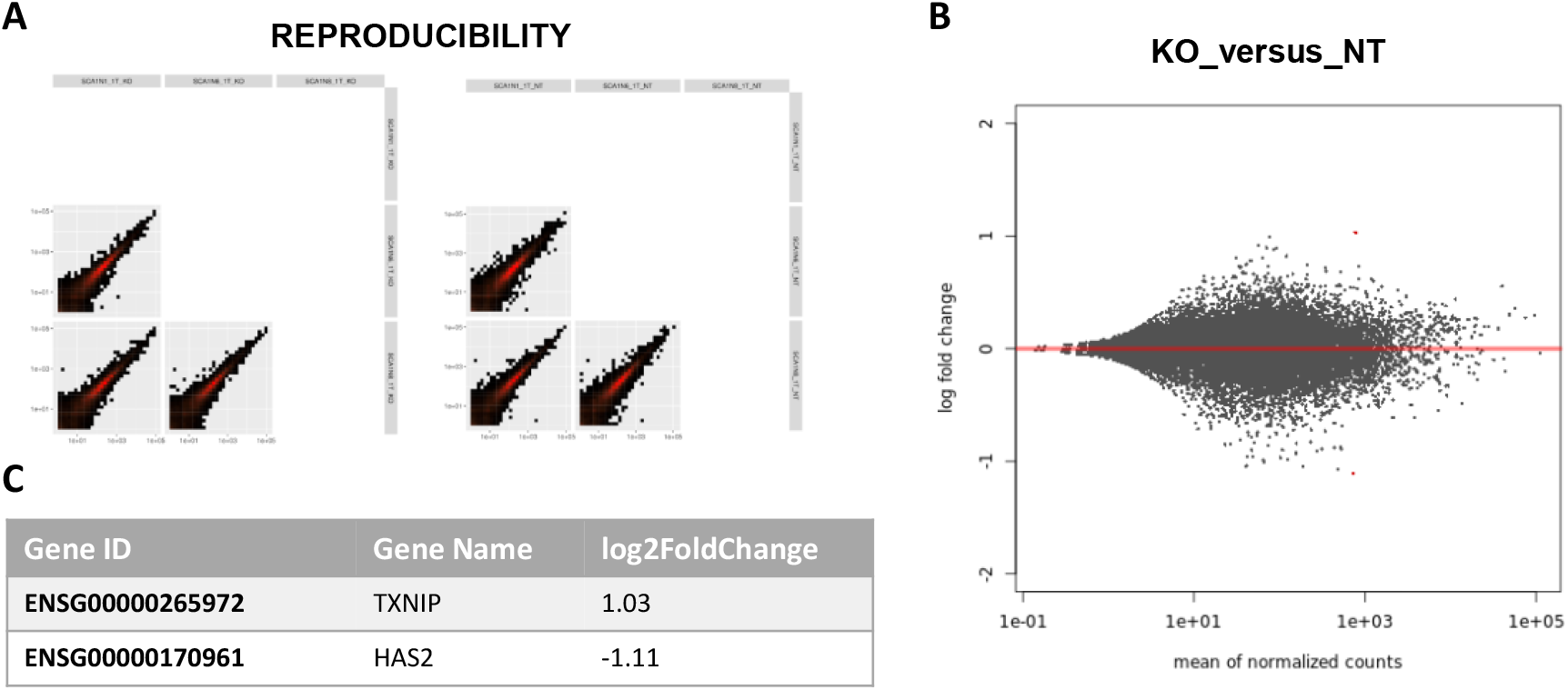
Trascriptome profile by RNA-seq. **A**, Inter-replicate correlation plots, which verify the overall correlation of expression between replicates. **B**, Digital gene expression data can be visualized as MA plot, just as with microarray data where each dot represents a gene. This plot shows RNA-seq gene expression for sgRNA/Cas9 treated versus not treated SCA1 fibroblasts. The red dots correspond to the only two differentially expressed genes, following treatment, showing in table **C**.

## Discussion

SCA1 is one of the two spinocerebellar ataxias with the highest incidence in Italy, especially in the North, with a frequency of around 21% [Brusco et al., 2004]. It is also present with a high relative prevalence in Russia, South Africa, Serbia and India [Di Donato et al., 2012]. There is no cure for SCA1, and current therapies only provide symptomatic relief. Although several research teams are trying to develop therapeutic strategies that can silence the *ATXN1* gene and block the production of the toxic protein, none of these approaches have entered clinical trials, neither in Europe [https://www.clinicaltrialsregister.eu/] nor in the United States [https://clinicaltrials.gov].

Recent studies have shown that polyQ-expanded ATXN1 exerts cerebellar toxicity mainly through its interaction with CIC [Rousseaux et al., 2018], therefore, genetic approaches aimed at blocking the production of this protein seem to be the most promising strategies for the treatment of SCA1. Since it has already been shown that specific CRISPR/Cas9-mediated gene editing could be used to permanently eliminate polyglutamine expansion-mediated neuronal toxicity [Ouyang et al., 2018; Yang et al., 2017], we successfully developed a CRISPR/Cas9-based approach to efficiently reduce the production of both healthy and mutated ATXN1 protein.

This therapeutic genome-editing strategy is not able to discriminate between the mutated and wild type allele of the *ATXN1* gene, but in any case, it has already been shown that a partial suppression of both forms of the ATXN1 protein is well tolerated [Keiser et al., 2016].

Our approach involves the use of two different sgRNAs targeting exon 8 of the *ATXN1* gene [Maeder and Gersbach, 2016] and, analysing the gene modifications introduced by our CRISPR/Cas9 system to the *ATXN1* gene, we found that the two sgRNAs seem to have different efficiency to recognise the target sequence and mediate the Cas9 cut. G3 and G8 sgRNAs mediate the excision of the polyQ tract only very few times (only 5% in SCA1N8), probably due to the steric hindrance caused by the nearness of the target sequences and the high molecular weight of the Cas9. G3 sgRNA showed a higher efficacy (with a ratio of about 3 to 1) and the largest number of indel and/or point mutations were localized upstream the polyQ tract compared to the cut mediated by G8 sgRNA downstream of the polyQ. Moreover, to verify if the lower efficiency of G8 was due to a guide design error, we analysed the theoretical efficiency of this sgRNA using all the software currently available for RNA design guides [Brazelton et al., 2015]. This analysis demonstrated the good theoretical efficiency of the G8 confirming our hypothesis that, in association with G3, there is a competition between the two guides and the dimensions of Cas9 which reduce the effectiveness of the G8. Therefore, we are designing a new RNA guide capable of recognizing a sequence much further downstream than that recognized by G8, in order to minimize the steric hindrance of the two sgRNA/Cas9 complexes. Although the CRISPR/Cas9 system has a minimal incidence of off-target effects [Kadam et al., 2018], variable levels of the latter have been observed [Fu et al., 2013; Hsu et al., 2013; Lin et al., 2014]. The Cas9 endonucleases produced in recent times are engineered to increase their efficiency towards the target sequence, minimizing the off-target effects. Furthermore, accurate design of sgRNAs and the direct delivery of the ribonucleoprotein complex, whose short half-life considerably reduces the exposure time of the cell genome to the action of the CRISPR/Cas9 system, can significantly reduce any off-target effects [Brazelton et al., 2015]. From the RNA-seq analysis, performed on three cell samples treated with G3/Cas9 and G8/Cas9 RNPs, we identified two genes whose expression was significantly, albeit in a limited way, altered by treatment: *HAS2*, downregulated, and *TXNIP*, upregulated. Since none of these are reported as possible off-site targeting of our system, we hypothesized that their variation is due to the suppression of the mutated ATXN1 protein and its toxic functions. Further studies will be done to investigate this aspect and to confirm that our silencing system can restore wild type conditions at the level of protein expression.

In conclusion, in this study we demonstrate to successfully reduce protein expression from SCA1 patient-specific cells using the CRISPR/Cas9 system, obtaining suppression efficiencies that agree with those obtained by Friedrich et al. when treated Atxn154Q/2Q mice with the antisense oligonucleotide ATXN1 ASO353 [Friedrich et al., 2018], but with the advantage of permanent suppression, without the need of continuous administration. However, these results will be confirmed in SCA1 fibroblasts of other patients that have been already recruited.

The CRISPR/Cas9 system is proving to be an effective and powerful technique for modifying genes. The use of the gene editing system in dominant inherited ataxias like SCA1, where the polyQ expansion in exon 8 of *ATXN1* leads to a gain of toxic protein function, may be the only solution for these neurodegenerative diseases.

## Materials and Methods

### sgRNA design and screening

sgRNAs were designed by CRISPR Design software, developed by Zhang at the MIT Laboratory in 2015 [Brezelton et al., 2015]. To test the efficacy of designed sgRNA, Guide-it In Vitro Transcription and Screening Kit (Takara Bio USA, Mountain View, California) was used, according to the manufacturer’s protocol.

### Sample-size estimation

The calculation of the sample size was made, assuming a 30% variation in efficacy in the treatment compared to the negative control, a 1st type error α equal to 0.05 and a power β of 0.80. The sample size calculation was equal to 5 cell samples.

### Skin biopsy

Patients underwent skin biopsy at the distal leg, 10 cm above the lateral malleolus, using a disposable 4-mm punch under sterile condition after local anaesthesia with lidocaine. The procedure does not need suture. Specimens were transferred into 15 ml tubes containing DMEM high glucose medium supplemented with 20% fetal bovine serum (FBS), antibiotic and antimycotic solution, and then taken to the laboratory to be processed. All subjects signed the informed consent and the studies were approved by the AVEC ethics committee.

### Fibroblasts isolation

The skin fragments were transferred to a 10 cm diameter cell plate and further fragmented with a disposable scalpel. The skin fragments were then moved into two T25 flasks with 1 ml of medium. Fresh medium was added after 48 hours and at regular intervals until it reached 3 - 4 ml in total, after which it was replaced twice a week. After 20-25 days, fibroblasts were detached with Trypsin-EDTA 1X (EuroClone, Pero, Italy) and moved to a new flask. New medium was added to the plate with skin fragments and fibroblasts were periodically transferred to new flasks for at least a month. Once amplified, SCA1 patient-derived fibroblasts were frozen as passage 1.

### Cell culture and transfection

SCA1 skin fibroblasts were maintained in DMEM high glucose medium supplemented with 20% FBS, antibiotic and antimycotic solution. 80 pmoles of Cas9 were complexed with 240 pmoles of sgRNA, incubating at room temperature (RT) for 20 minutes. RNP complexes were transfected in 500,000 cells with Amaxa Nucleofector II (Amaxa Biosystems, Lonza, Basel Switzerland) using the P-022 program according to manufacturer’s protocol and adding 1 ul of 100 μM Alt-R^®^ Cas9 Electroporation Enhancer (IDT, Coralville, Iowa), followed by plating in 6 well plates. The cells were then amplified for at least 7-10 days until a sufficient number of cells was obtained, from which both genomic DNA and total proteins were extracted.

### Western blotting analyses

Protein extracts were prepared by homogenization of cellular precipitates in extraction buffer (50 mM Tris-HCl pH 8.0, 1% V/V Triton X-100, 0.5% V/V Nonidet P-40, 10 mM mercaptoethanol, 4% V/V glycerol and Complete Mini Protease inhibitor cocktail tablets (Roche, Basel Switzerland), 1 tablet in 10 mL), followed by five freeze/thaw cycles and then centrifuged at 4°C for 3 min at 14,000 rpm. Supernatants were quantified by Pierce™ BCA Protein Assay Kit (Thermo Fisher Scientific, Waltham, Massachusetts) and used for Western blotting analysis, using for the ATXN1 detection the antiserum 12NQ, kindly donated by Dr. Orr [Perez Ortiz et al., 2018], and the antibody against HSP90 (Biosciences Inc., Allentown, Pennsylvania), as housekeeping. 30 μg of protein extracts were denatured for 5 minutes at 98°C in 1X SDS sample buffer (62.5 mM Tris-HCl pH 6.8, 2% SDS, 50 mM Dithiotreithol (DTT), 0.01 % bromophenol blue, 10% glicerol), resolved by SDS-PAGE and transferred to nitrocellulose membranes (BioRad Laboratories Inc., Hercules, California). After blocking with 3% skim milk/TBS, the membranes were incubated with primary antibodies in 1% skim milk/TBS overnight at 4°C. After washing in 0.05% Tween 20/TBS, membranes were incubated with the corresponding secondary antibody conjugated with HRP in 1% skim milk/TBS for 1 hr at RT and washed again. Chemiluminescence signals were detected using the Clarity Western ECL Substrate (BioRad Laboratories Inc.) according to the manufacturer’s protocol. The signal intensity was determined using the ChemiDoc MP Imaging System (BioRad Laboratories Inc.) and ATXN1 expression in treated cells compared to not treated ones was calculated using for normalization both total proteins amount and HSP-90.

### Phenol/chloroform extraction of genomic DNA

Cell pellets were resuspended with a lysis solution (10 mM Tris HCl pH 8.0, 400 mM NaCl, 2 mM EDTA pH 8.0, 0.45% p/V SDS and 0.45 mg/mL proteinase K) and incubated at 56°C for 2 hours. One volume of phenol:chloroform:isoamyl alcohol (25:24:1) was added and mixed by inversion for 5 minutes. Samples were centrifuged at RT for 5 minutes at 12,000 rpm and then the aqueous phase was transferred to a clean tube. After a second extraction to optimize DNA purification, an equal volume of chloroform:isoamyl alcohol was added to the aqueous phase and samples were centrifuged under the same conditions described above. 2.5 volumes of 96% ethanol were added to each tube, which were placed at –20°C overnight to precipitate DNA. Samples were centrifuged at 4°C for 5 minutes at 12,000 rpm, washed with 1 ml of 70% ethanol and then centrifuged again under the same conditions. The dry pellets were finally resuspended in water and the DNA quantified using the Biospectrometer (Eppendorf, Hamburg, Germany).

### PCR

PCR products were amplified in 100 ml reactions with 0.2 units of EconoTaq DNA polymerase (Lucigen, Middleton, Wisconsin), 1 μM of each primer, and 0.25 mM of each dNTP. Thermal cycling was done using an annealing temperature of 56°C. PCR products were purified using NucleoSpin^®^ Gel and PCR Clean-up (Macherey-Nagel, Hoerdt, France), according to manufacturer’s protocol.

### PCR subcloning

Purified PCR fragments were cloned into a pGEM-T Easy Vector (Promega, Madison, Wisconsin), according to the manufacturer’s protocol. The ligation reactions were then transformed into E. coli 5-alpha Chemically Competent cells (Lucigen), performing a thermal shock for 45 seconds at 42°C followed by 2 minutes on ice. After 1 hours at 37°C in slow agitation, bacteria were plated in 100 ng/μl ampicillin LB-Agar plates, with 0.01% X-gal/200 μM IPTG.

White colonies were amplified and plasmids purified using PureYield™ Plasmid Miniprep System (Promega), according to the manufacturer’s protocol. The obtained plasmids were analysed through electrophoresis in 0.8% agarose gel, quantified using the Biospectrometer (Eppendorf) and send to BMR Genomics for sequencing, set up using the M13 forward sequencing primer.

### Total RNA extraction

The total cellular RNA was extracted by TRIZOL^®^ Reagent (Gibco - Thermo Fisher Scientific), according to manufacturer’s protocol. All reagents and materials used were RNase-free. The extracted RNA was analysed by electrophoresis on 0.8% agarose gel and sent to Lexogen (Greenland, New Hampshire) for RNA-seq analyses.

### Statistical analysis

Statistical analyses were performed using GraphPad Prism 7.00 software. The statistical tests applied were: unpaired t test with two-tailed P value and alpha level P<0.05; F test to compare variances with alpha level P<0.05.

## Abbreviations

SCA1: Spinocerebellar Ataxia type 1
ATXN1: ataxin 1
PolyQ: polyglutammine
ASO: antisense oligonucleotide
sgRNA: single guide RNA
CRISPR: clustered regularly interspaced short palindromic repeat
Cas: CRISPR associated proteins
RNP: ribonucleoprotein

## Acknowledgements

We thank Harry T. Orr for providing us with anti-ATXN1 antiserum 12NQ for our western blotting experiments.

